# scCorr: A graph-based k-partitioning approach for single-cell gene-gene correlation analysis

**DOI:** 10.1101/2021.03.04.433945

**Authors:** Heng Xu, Ying Hu, Xinyu Zhang, Bradley E. Aouizerat, Chunhua Yan, Ke Xu

**Author notes:** Correspondence: Ke Xu, MD, PhD, Department of Psychiatry, Yale School of Medicine, 950 Campbell Ave, Building 36, West Haven, CT 06511. contributed equally (Heng Xu, Ying Hu).

## Abstract

An important challenge in single-cell RNA-sequencing analysis is the abundance of zero values, which results in biased estimation of gene-gene correlations for downstream analyses. Here, we present a novel graph-based k-partitioning method by merging “homology” cells to reduce the number of zero values. Our method is robust and reliable for the detection of correlated gene pairs, which is fundamental to network construction, gene-gene interaction, and cellular -omic analyses.

## Main

Single-cell RNA-sequencing (scRNA-seq) enables transcriptome profiling at high cell resolution and provides unprecedented precision into the molecular mechanisms underlying disease^1,2^. A central challenge to cell type identification and downstream analysis is the abundance of zero values, known as “dropout”, in single cells due to either low transcript copy number and/or ineffective capture capacity of scRNA-seq technology. scRNA-seq typically captures only 5-15% of the transcriptome of each cell^3^. Dropout causes significant zero inflation, increasing background noise, which leads to loss of detection of gene-gene correlations crucial to gene network construction and determination of lineage relationships among cells.

Considerable computational effort has been expended to address scRNA-seq dropout. One approach is to aggregate cells using small proportions of highly variable genes^4^ that are heavily weighted in analysis. Another approach is to impute zero values^5–7^. For example, DeepImpute employs a deep neural network imputation algorithm that uses dropout layers to identify patterns in scRNA-seq data to impute zero values^8^. Markov Affinity-based Graph Imputation of Cells imputes likely missing expression data to detect the underlying biological structure via data diffusion^9^. A newly developed algorithm embraces zero values based on binary zero/nonzero patterns into the analysis to improve gene-gene correlation and gene network analysis^10^. While these methods reduce dropout, recovering gene-gene relationships from zero abundant data remains challenging due to the noise introduced by imputation of a large number zero values or loss of information by simplifying the complexity of data.

Here we present “scCorr”, a novel graph-based k-partitioning approach to address dropout by recovering missing gene-gene correlations. The scCorr algorithm: 1) generates a graph or topological structure of cells in scRNA-seq data; 2) partitions the graph into k multiple min-clusters employing the Louvain algorithm, with cells in each cluster being approximately homologous (with similar transcriptional profiles); 3) visualizes the series of k-partition results to determine the number of clusters; 4) averages the expression values, including zero values, for each gene within a cluster; and 5) estimates gene-gene correlations within a partitioned cluster. We demonstrate that the graph k-partitioning approach enables the reliable acquisition of cluster-based gene-gene correlations in a peripheral blood mononuclear cell (PMBCs) scRNA-seq dataset. We applied scCorr to estimate the accuracy of cell type identification in a known cell type (i.e. CD4^+^ T cell) in the dataset and compared the performance of gene-gene correlation estimation between scCorr and conventional non-clustering single-cell gene correlation methods.

We first confirmed a significant abundance of zero values in the data generated from 15,973 PBMCs^11^. A total of 21,430 genes were annotated; we observed that 95% of the 21,430 genes had zero values in a cell (**Figure 1A**), and 95% of the 15,973 cells showed at least one undetected gene with zero values (**Figure 1B**), suggesting that scRNA-seq captures only 5% of gene expression at the single cell level. For example, among the 347 genes that compose 1,532 biologically related gene pairs on 3 Kyoto Encyclopedia of Genes and Genomes (KEGG) pathways (i.e. hsa04010, hsa04115, hsa04662) (**Table S1**), over 70% exhibited zero expression (**Figure 1C**, **1D**), resulting in poor gene-gene correlation among these known gene pairs. These data underscore the importance of developing a method to minimize zero value inflation to recover gene-gene correlations.

**Figure 1.**
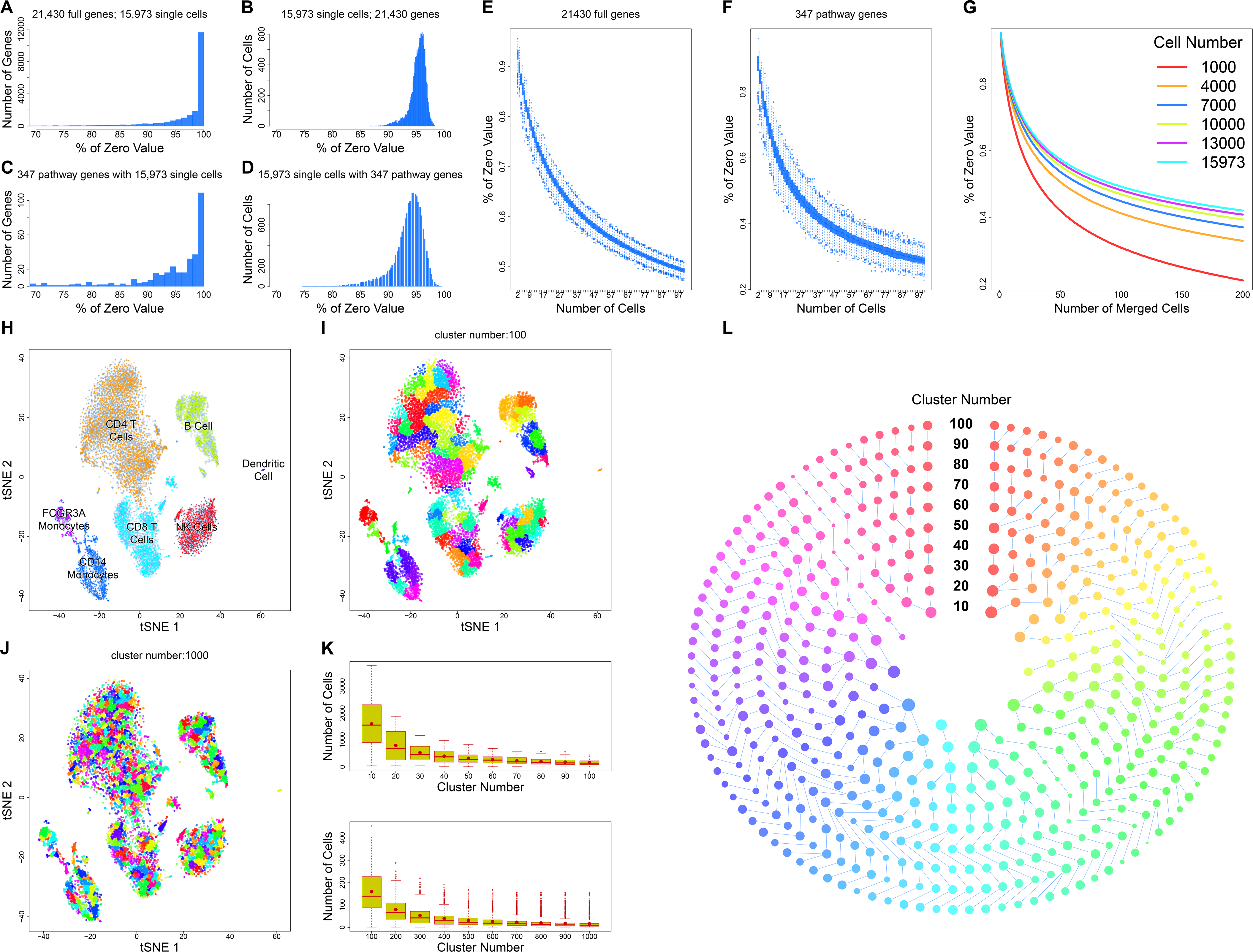
Study principals and workflow. Data are from single-cell RNA-seq of peripheral blood monocyte cells (PBMCs) in healthy participants. **A**-**D** present the distribution of zero values by individual genes and by single cells. A total of 21,430 genes had zero values in at least one cell (**A**), and more than 95% of 15,973 cells showed zero values in at least one cell (**B**). Among a set of 347 genes from 3 KEGG pathways, all genes had zero values in at least one cell (**C**), and 95% of 15,973 cells contained zero values in at least one gene (**D**). **E**-**G** show reductions of zero values in merged cells. The percentage of zero values of 21,430 genes was markedly reduced in merged cells. The zero-value reduction is approximately 50% among 50 merged cells (**E**). Similarly, zero values of the 347 genes were reduced in merged cells (**F**) and was consistently observed in 6 different cell sets (**G**). **H**-**L** present the workflow and features of the scCorr method. First, data dimensional reduction and cell classification by tSNE and cell type identification using the marker gene approach (**H**). Second, cell partitioning was based on a tSNE plot by using scCorr with different numbers of clusters (**I**: k=100; **J**: k=1,000). Average number of cells per cluster is shown. The red dot is an expected number of cells in a cluster (**K**). ScCorr enables tracing of the evolution of each partitioned cluster (**L**).

We observed that merging transcriptomically homologous cells dramatically reduced the proportion of zero values. In a set of 50 merged cells that were randomly selected from the 15,973 PBMCs, the percentage of zero values from the same set of 21,430 genes was reduced from 90% in two to 57.4% (95% confidence interval [CI), 57.3%, 57.4%] in the 50 merged cells (**Figure 1E**). Revisiting the same 347 genes from the 3 KEGG pathways, the reduction in zero values increased further, from 90% in two to 37.6 % (95% CI: 37.5%, 37.8%) in the same 50 merged cells (**Figure 1F**). Examining the impact of the number of merged cells on zero value frequency, focusing on six cell populations of 1,000 to 15,973 cells, the reduction of zero value proportion stabilized when the merged cells exceeded 100 in a set of 1,000 cells (**Figure 1G**). Simulation analysis of four separate sets of cells and genes further supported the reduction of zero values in merged cells (**Supplementary Figure 1A** and **1B**). This property motivated the development of the scCorr package.

scCorr is based on local optimal modularity (i.e. the Louvain algorithm) to partition a graph into k clusters. **Figure 1H-1K** presents the analytic strategy of scCorr on the scRNA-seq data (21,430 genes, 15,973 cells), for which tSNE identifies seven major cell types (**Figure 1H**). The graph edges are weighted by the distances and distance matrix, which is converted to a weighted graph determined by a cutoff of three as the default (any edge weight > 3 is removed from the graph), and an initial number of N clusters is generated with N greater than k. Next, the center of each cluster is calculated, and adjacent clusters with the smallest distance are merged one by one until N equals k. As presented in **Figure 1I** (k=100) and **1J** (k=1,000), scCorr-partitioned clusters do not change the local structure of transcriptomically-defined cell populations. A series partitioning process was carried out to determine the desired number of clusters (**Supplementary Figure 2**). **Figure 1K** displays the cell number distributions for each cluster set. For each box plot, the observed number of cells for each cluster (median) is close to an expected number of cells per cluster. One of the utilities included in the scCorr package is to trace how a cluster evolves during k graph partitioning in multiple cluster sets, displaying ladder and circle plots to visualize the evolutionary process of each cell cluster in a variety of cluster sizes (**Supplementary Figure 3**). As an example, **Figure 1L** depicts the evolutionary process for each cluster during graph partitioning.

Next, based on averaged gene expression values in a partitioned cluster we computed gene-gene correlations using a generalized linear model. We estimated gene-gene correlations in the CD4^+^ T cell population, focusing on the same 347 genes and 1,532 gene pairs defined by the 3 KEGG pathways. In this instance, we present the results of scCorr-partitioned cluster sizes from 40 to 1,000, which shows greater gene-gene correlation coefficients and identifies more gene pairs than the conventional noncluster single-cell-based method. For example, in a set of 100 clusters, scCorr and the noncluster method identified a combined total of 85 of 1,242 possible gene pairs (71 solely by scCorr, 10 solely by the noncluster single-cell method) with only 4 detected by both methods (false discovery rate (FDR)<0.05). Gene-gene correlations were stronger among the pairs detected by scCorr than by the noncluster single-cell method (62 of 85 [73%] of the gene pairs with r>0.5 were detected by scCorr, while 4 of 85 [5%] had r>0.1 using the noncluster single cell-based method). Among significant gene pairs detected by both methods, approximately 50% of gene-gene correlations were in the agreement (blue dot) and the other 50% in disagreement (red dot) in the direction of the correlations (**Figure 2**). As an example, **Figure 2A** and **2B** presents the p-values and r-values of the top 45 gene pairs selected by p-value (scCorr: 40, noncluster single cell-based method: 14, 9 overlapping). For example, *ATF4* and *MAPKAPK2* are a well-established pair of co-expressed genes. scCorr increased the coexpression of *ATF4* and *MAPKAPK2* (r=0.82) (**Figure 2C**) compared to the noncluster gene-gene correlation method (r=0.01) (**Figure 2D**). Similarly, scCorr outperformed the correlation of another co-expressed gene pair, *MAPK1 and DUSP2* (scCorr: r=0.586, noncluster single cell-based method: r=0.007) (**Supplementary Figure 4**). Of note, *scCorr detected no significant correlations for randomly selected genes*. These results suggest that scCorr shows robust detection of correlated gene pairs by minimizing zero-value effects.

**Figure 2.**
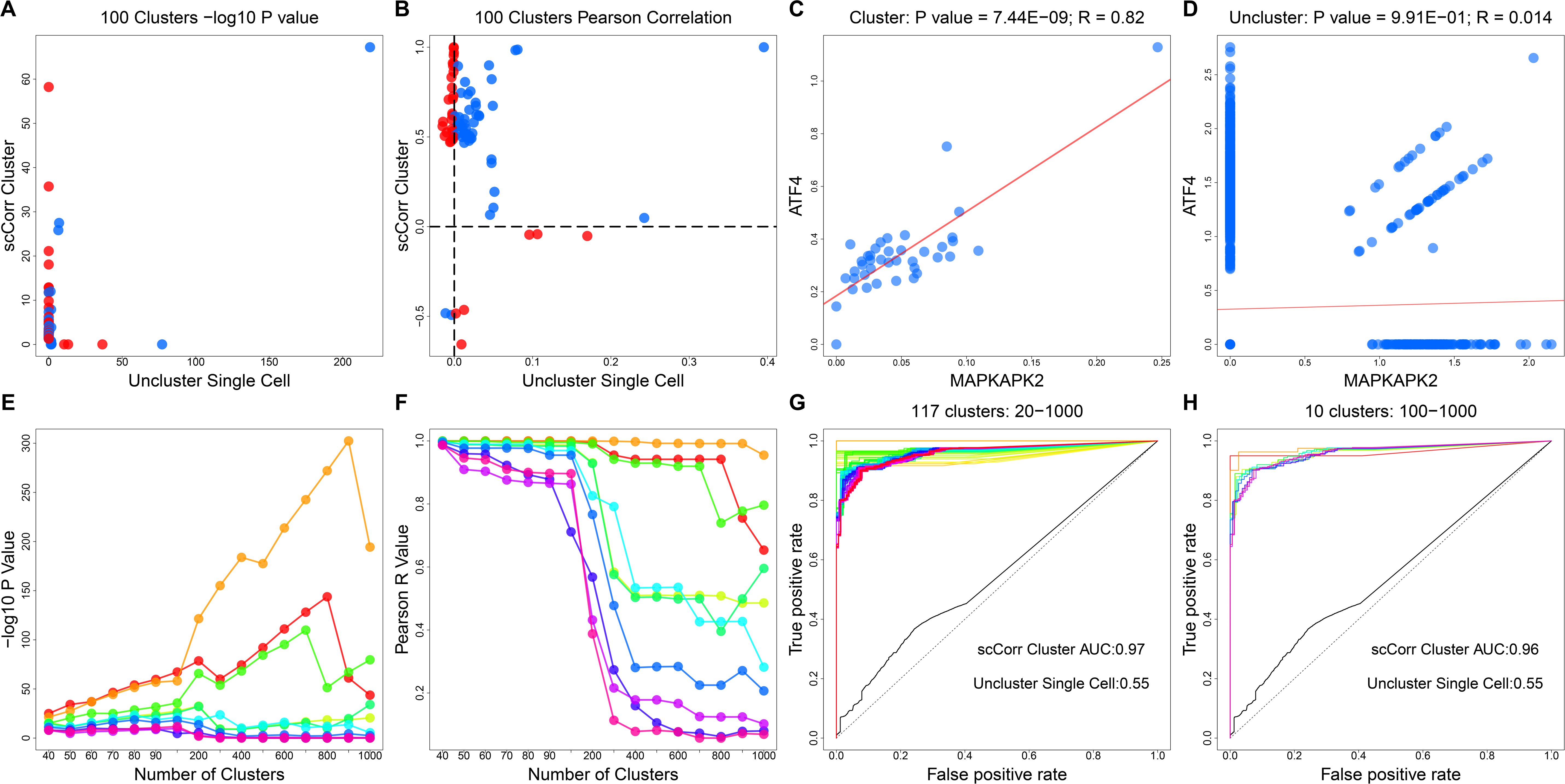
Evaluations of scCorr performance in gene-gene correlations among CD4+ T cells (**A**-**F**) and cell type identification (**G**-**H**). Gene-gene relationships were quantified by Pearson correlation. In **A** and **B**, gene-gene correlation was separately performed in nonclustered single cells (X-axis) and in scCorr-partitioned cell clusters (K=100) (Y-axis). Only the top 10 significantly correlated gene pairs are shown. Red dots indicate agreement and blue dots indicate opposite directions of gene-gene correlation between two methods. Correlated genes are shown as –log_10_ p- (**A**) and r- (**B**) values. scCorr detected significant correlation between MAPKAPK2 and ATF4 genes (**C**) while the conventional noncluster single cell-based method showed no significant correlation between the two genes (**D**). Gene-gene correlation varies in different numbers of clusters. **E** and **F** show the top 10 correlated genes in different numbers of clusters partitioned by scCorr among CD4+ T cells evaluated by –log_10_ p- (**E**) and r- (**F**) value. The performance of scCorr for cell type identification of CD4+ T cells are shown in **G** (k=117) and **H** (k=10). The Receiver Operator Characteristic Area Under the Curve (AUC) was greater when using scCorr (AUC=0.97 and 0.96) than when using nonclustered single cells (AUC=0.55).

One key step in scCorr is to establish a reasonable cluster size cutoff. We recommend performing a series of correlation analyses with different cluster sizes in order to identify the cluster size at which stable p- and r-values in the cluster sets are achieved (**Supplementary Figure 5**). For example, in a set of the top 10 most significant gene pairs in a series of cluster sizes from 40 to 1,000 in CD4^+^ T cells, the p- and r-values were relatively stable in cluster sizes from 40 to 100 (**Figure 2E and 2F**). However, gene correlation changed dramatically when the cluster size exceeded 100. As a result, a size of 100 clusters each containing 125 homologous cells showed the greatest correlation coefficients and smallest p-values in the dataset, indicating that a cluster size of 100 is a reasonable cutoff to achieve consistent gene-gene correlations in CD4^+^ T cells.

Finally, we tested the efficiency of scCorr in predicting cell type classification. We performed 10-fold cross-validation to predict CD4^+^ T cells in our dataset^11^. The average of the Receiver Operator Characteristic area under the curve (AUC) was 0.97 across 117 (**Figure 2G**) and 0.96 across 10 (**Figure 2H**) clusters partitioned by scCorr. In contrast, the performance of noncluster single cell-based prediction was poor (i.e., AUC=0.55) from the same number of nonclustered cells, suggesting that scCorr-based cell type identification is reliable, accurate and outperforms the noncluster single cell-based approach.

scCorr requires a reasonable amount of computational time. Users are able to adjust the computational time by estimating a range of scales and k partitioning clusters (**Supplementary Figure 6A**). In a set of 5,967 cells, the estimated computation time for a scale of 200 is approximately 10 minutes (**Supplementary Figure 6B**). The computation time for a scale of 400 is estimated to be 20 minutes for a set of 15,973 cells (**Supplementary Figure 6C**).

In summary, built on the observation that merging homologous cells can decrease the number of zero values in a given cell type, scCorr was developed based on a graphic structure of cells in homologous cells to estimate gene expression values, including zero values, in a local cell community. Unlike imputation methods, which infer missing gene expression, possibly introducing noise into the data, scCorr maintains local community data organization in single cells by k-partitioning a graph into mini-clusters in the similar cell-cell matrix based on transcriptomic similarities within the cell type. Thus, the merged cells in a partitioned cluster increase the power to recover correlated gene pairs. We show that cluster-based analysis by scCorr k-partitioning detected more significant gene pairs than noncluster single cell-based analysis, suggesting that scCorr is a robust approach to address dropout in scRNA-seq. scCorr is robust, simple, fast, and easy to use in the R environment. The method can be broadly applied to single-cell genomic analysis and is particularly useful for downstream analyses, such as differential gene expression analysis, gene-set enrichment analysis, and network construction at single-cell resolution.

## Methods

### Single-cell RNA-seq data

Single-cell data were downloaded from the NCBI Gene Expression Omnibus (GEO) https://www.ncbi.nlm.nih.gov/geo/query/acc.cgi?acc=GSE130228. The dataset was generated using the 10x Genomics platform and contains 15,973 single cells, 21,430 genes and seven cell types. Single-cell data quality control, normalization, and cell type identification were described previously^11^.

### Correlated genes in KEGG pathways

A total of 1,532 gene-gene interactions from 347 unique genes were selected from the B cell receptor, p53, and MAPK signaling pathways in the KEGG database (https://www.genome.jp/kegg/pathway.html) (**Supplementary Table 2**). The 1,532 gene pairs served as a reference in the scCorr analysis using the scRNA-seq dataset described above.

### Zero value distribution in merged cells

The distribution of the percentage of zeros for each cell (**Figure 1B** and **1D**) and each gene (**Figure 1A** and **1C**) is shown using histogram plots. Two different gene sets were used: all 21,430 genes and only 347 genes on the 3 KEGG pathways from GSE130228. We examined the zero-value distribution in different sets of merged cells from GSE130228 and from a simulation dataset. We simulated 95% of zero values assuming 23,000 genes and 20,000 cells. We randomly selected two cells and calculated the percentage of zero values in 23,000 genes. Each two-cell selection was performed 1,000 times, and an average of zero percentage in 23,000 was determined. The selection of random cell numbers was repeated until 200 cells merged. We also simulated different numbers of cells (2,000, 8,000, and 14,000) and different sets of gene numbers (16,000, 18,000, and 20,000). The distributions of zero values were essentially the same (**Supplementary Figure 1**).

### Zero-value reduction through k-partitioning

The strategy to reduce zero values in gene expression involved using a k-partitioning algorithm to group cells into clusters and calculate the average gene expression in the cluster instead of in single cells. An adjacency distance matrix of single cells was estimated by the single-cell expression profile from the tSNE output and converted into a weighted graph or network using the R package igraph. The weight values were the length of the edges in the graph. The Louvain algorithm, an efficient graph-clustering method based on the modularity measure and a heuristic approach, was used to group cells into a predefined number of clusters through iteratively splitting and merging cell processes shown in the t-SNT plot of cluster numbers 50, 100 and 1000 (**Supplementary Figure 2A**). The pseudocodes are included at the end of the Methods section. Cell number distributions from cluster numbers 10-100 and 100-1000 are displayed using box plots (**Figure 1K**).

### Visualization of cell clusters using the k-partitioning algorithm

Multiple cluster visualization functions were implemented to evaluate the k-partitioning algorithm performance to select desired cluster numbers. In the cluster overlay on the t-SNE plot, each dot represents a cluster with dot size being proportional to cluster size (**Supplementary Figure 2B**). This provides a convenient function to identify uniformly distributed clusters. Using tree-based visualization of clusters, ladder and circle plots show the evolution of clusters at the different cluster numbers (**Supplementary Figure 3**). Dot size is proportional to the cell number in one cluster and lines between dots track the cluster development. **Supplementary Figure 3A** presents a tree of 20-40 clusters from top to bottom. The tree can be arranged in a circular shape suitable for displaying a large number of clusters (**Supplementary Figure 3B** and **3C**); inner and outer circles correspond to the top and bottom trees, respectively.

### Gene-gene correlation analysis

Pearson and Spearman correlation coefficients at single-cell and cluster levels were calculated between gene pairs extracted from KEGG pathways, as exemplified in the scatter and violin plots of MARPK1-DUSP2 (**Supplementary Figure 4**). The correlation p-values were estimated using generalized linear models (glm). At the cluster level, the average gene expression of cells in a cluster was used as the gene expression value. The top 10 correlated gene pairs by p-values based on 100 clusters were selected to evaluate the effect of the number of clusters on the correlation. A total of 16 cluster sets were used: 40-100 and 100-1000 in increments of 10 and 100, respectively. The relationships of glm p-values and Pearson and Spearman correlation coefficients vs different numbers of clusters are illustrated using a line plot (**Supplementary Figure 5**).

### Time estimation for k-partitioning clusters

We implemented calculation time for different numbers of cell clusters. In our analysis, using Rtsne with perplexity=30 and max_iter=2000, the x and y coordinate regions were approximately from −50 to 50. Before performing graph-based clustering, we suggest that the x and y coordinate regions are scaled from −200 to 200 or −400 to 400 in cell numbers 5,967 and 15,973, respectively. **Supplementary Figure 6** shows estimated times for a range of scaled clusters. On a tSNE plot scale of −50 to 50 of 5,976 cells, when the scale range is 200, the running time is smallest, regardless of the number of clusters. In a plot of 15,973 cells, a scale range of 400 appears to be the most rapid option, regardless of the number of clusters. The different colors for the lines represent the numbers of clusters.

## Codes

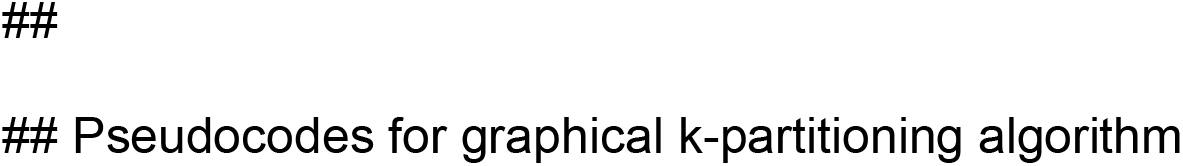
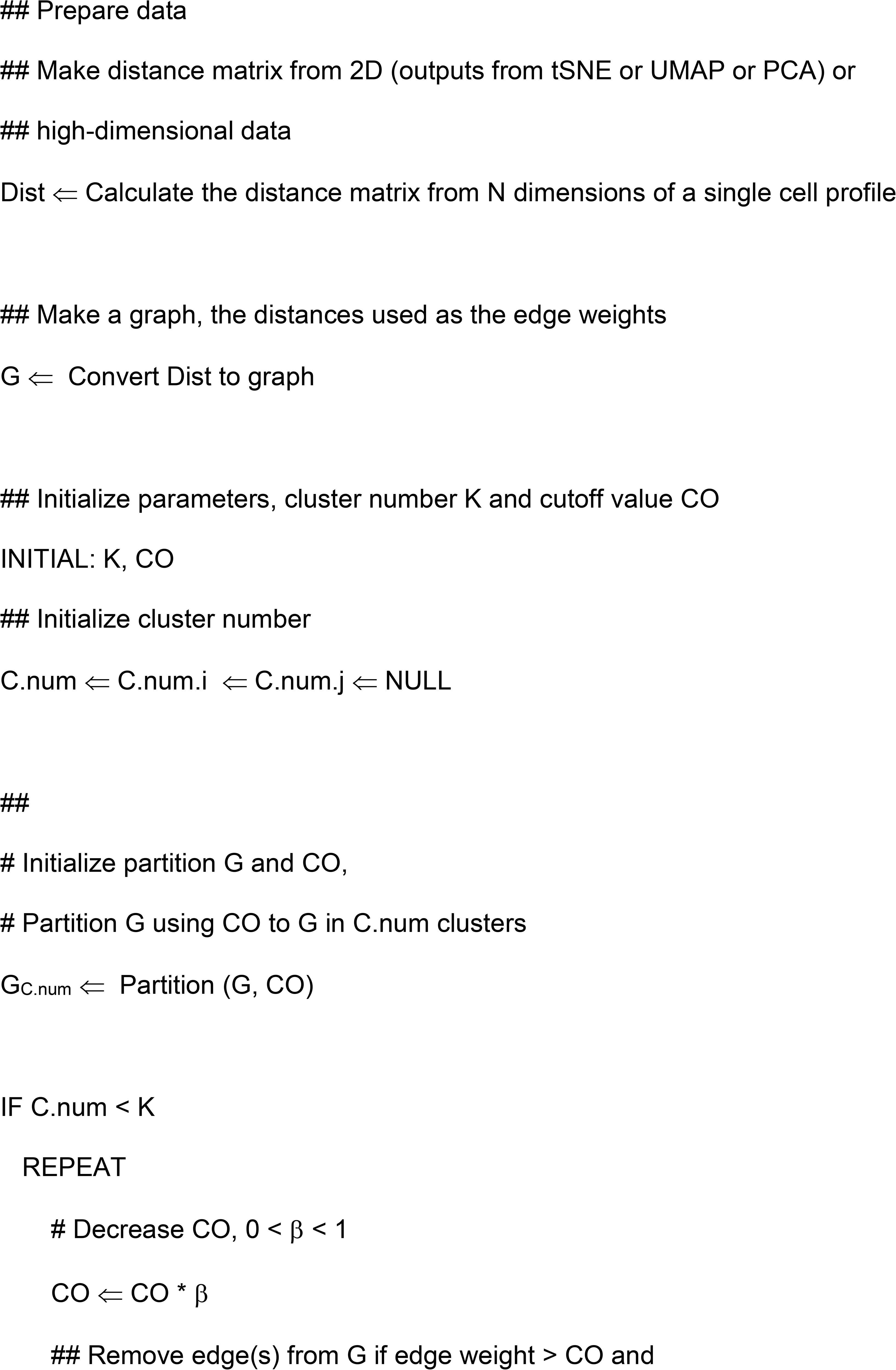
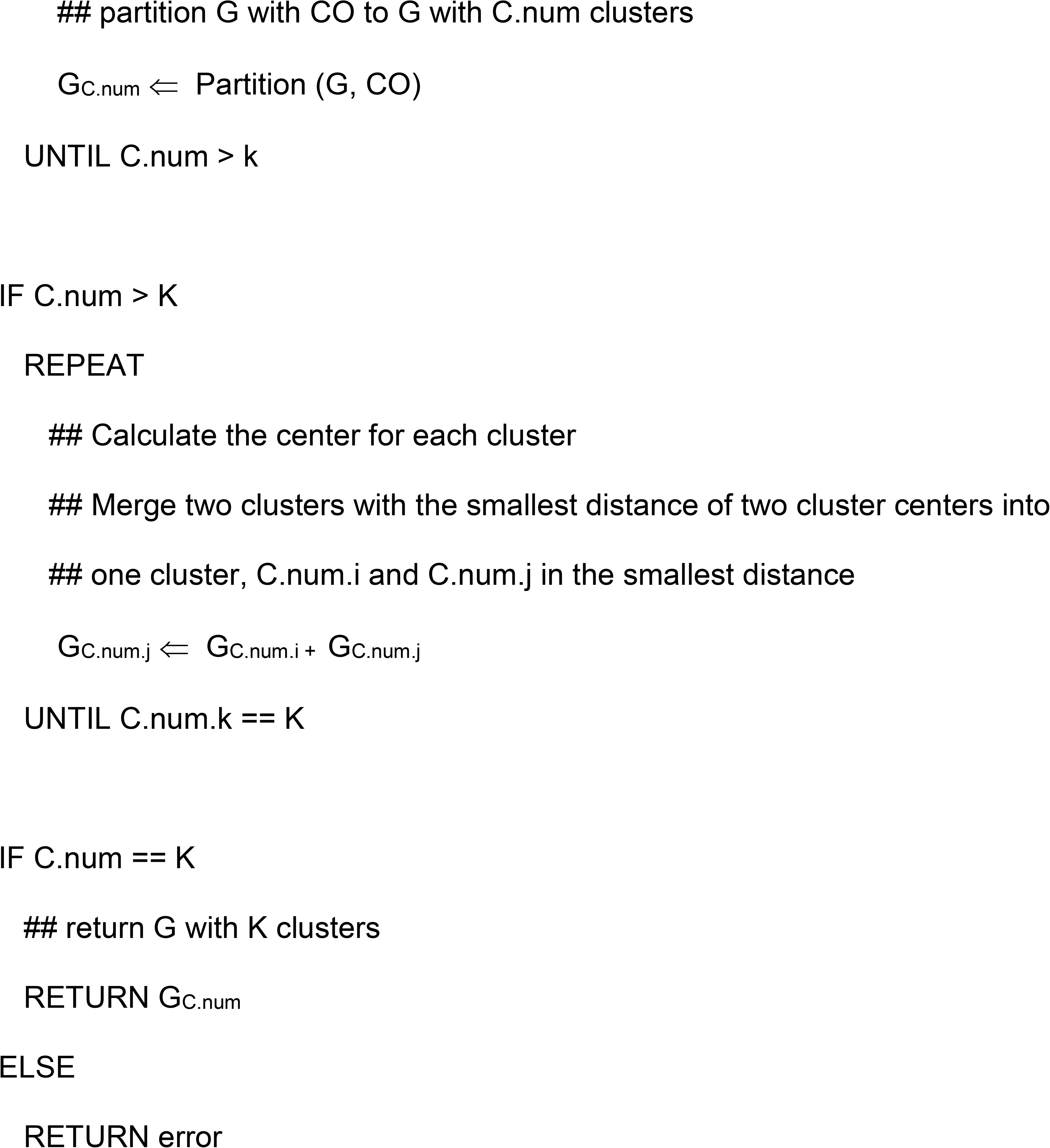

## Supporting information

Legends for Supplementary Figures

## Data availability

Sequencing data have been deposited to GEO under the accession GSE130228.

## Code availability

The scCorr source code is freely available from GitHub under the CBIIT-CGBB (https://github.com/CBIIT-CGBB/scCorr). The R package is also available from the Nature Research Reporting Summary.

## Contributions

H. X. and Y. H. carried out data analysis, coded the scCorr package, and prepared the first draft of the manuscript; X. Z. was involved in data analysis, B.E.A. was involved data presentation and manuscript preparation; C. Y. involved in the manuscript preparation; K. X. was responsible for study design, results interpretation, and manuscript preparation.

## Ethics declarations

The project was approved by Human Research Protection committee at Yale University.

## Competing interests

The authors declare no competing interests.

## Notes

### Competing Interest Statement

The authors have declared no competing interest.

