## Supplementary material for "scCorr: A graph-based k-partitioning approach for single-cell gene-gene correlation analysis": Legends for Supplementary Figures

*** contributed equally**

### Legends for Supplementary Figures

**Supp Figure 1**. Distributions of zero values of gene expression in four sets of simulated datasets (**A**) and in the scRNA-seq dataset with 21,430 genes and 15,973 peripheral blood mononuclear cells (**B**)

**Supp Figure 2**. t-SNE plot-based k-partitioning cluster. (**A**). All cells are clustered as the size of 50, 100, and 1,000 groups. (**B**). Three identical structure plots show a proportion of a cluster size. Each dot represents a cluster and the size of the dot is proportional to the cluster size.

**Supp Figure 3**. Tree-based visualization of cell clusters by the k-partitioning algorithm. Ladder clusters, N=20-40 (**A**); Circle clusters, N=20-40 (**B**); and Circle clusters, N=100-1,000 (**C**). The size of each dot represents the proportion of the cell number in one cluster.  A line connects two closest adjacent clusters.

**Supp Figure 4**. Correlation of a co-expression gene pair: DUSP2 and MAPK1 estimated by the non-clustering singal cell method (**A**) and by scCorr clustering method (**B**).

**Supp Figure 5**. Correlation of top 10 co-expressed gene pairs in different numbers of partitioned clusters in CD4+ T cells: evaluated by p values and r values using Person Correlation and Spearman Correlation.

**Supp Figure 6**. Estimation of computation time for tSNE-based k- partitioning cluster. xy.coordinate represents the regions of scaling. (A) Estimation of xy coordinates (B) In a tSNE plot scale of -50 to 50 of 5,976 cells, the running time is the shortest at the scale range is 200 regardless of the number of clusters. (C) In a plot of 15,973 cells, a scale range of 400 appears most rapid regardless of the number of clusters. Different color of line represents the numbers of clusters.
