## Supplementary figures and images for "scCorr: A graph-based k-partitioning approach for single-cell gene-gene correlation analysis"

### Supplemental Figure 1

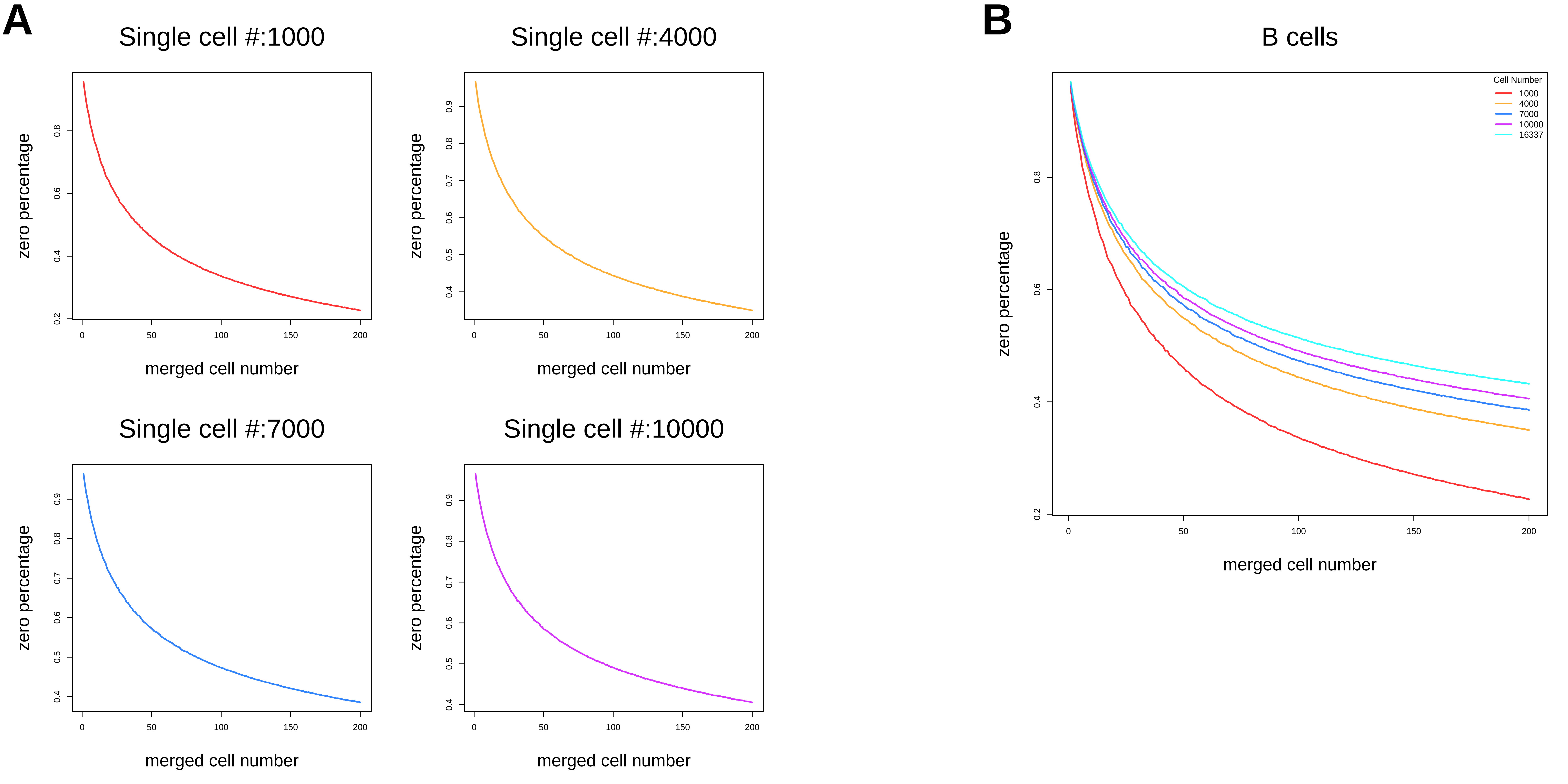

### Supplemental Figure 2

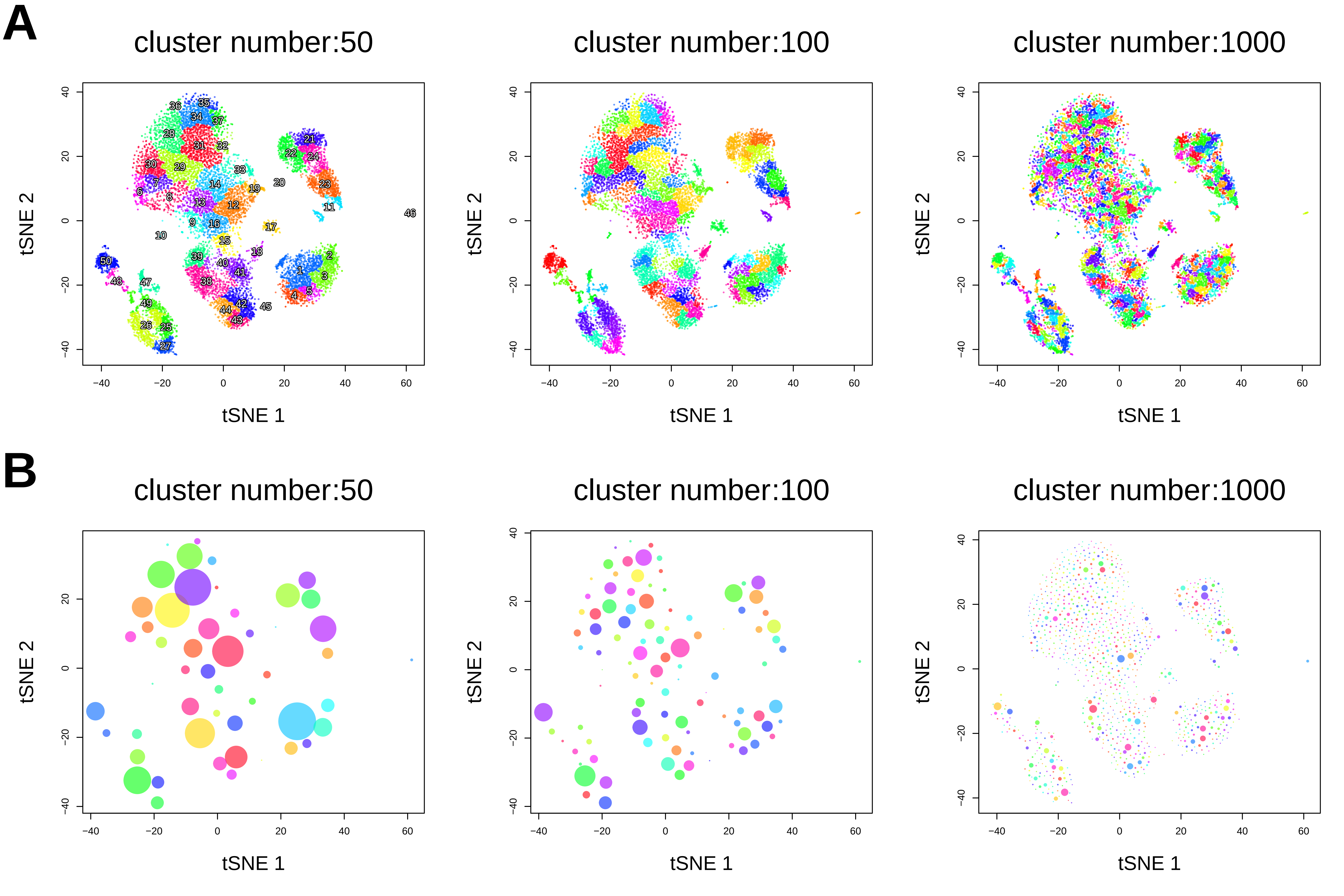

### Supplemental Figure 3

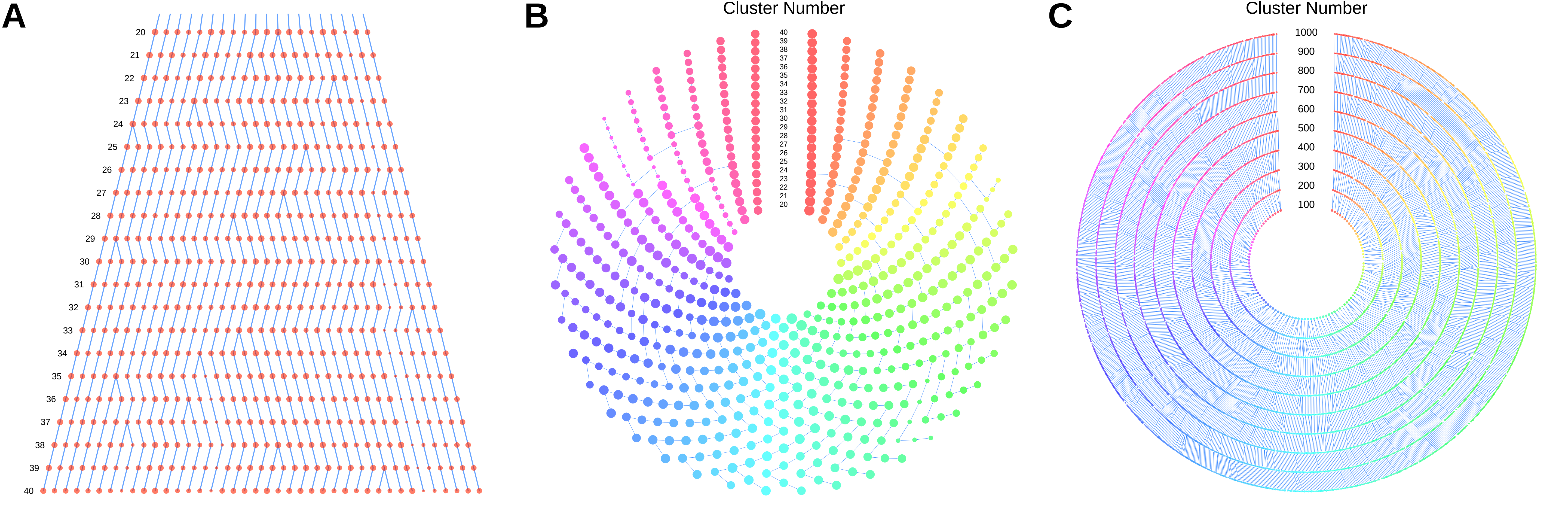

### Supplemental Figure 4

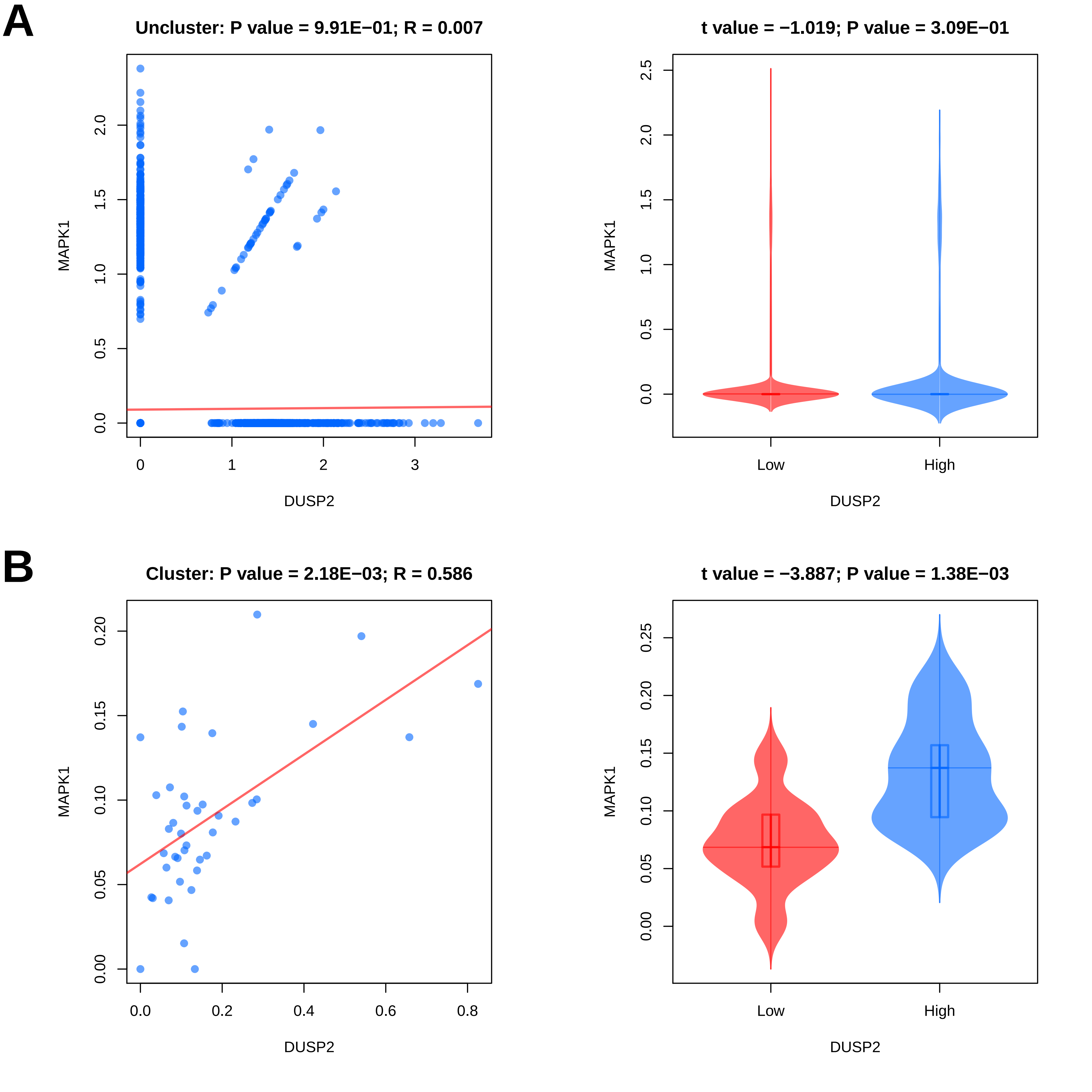

### Supplemental Figure 5

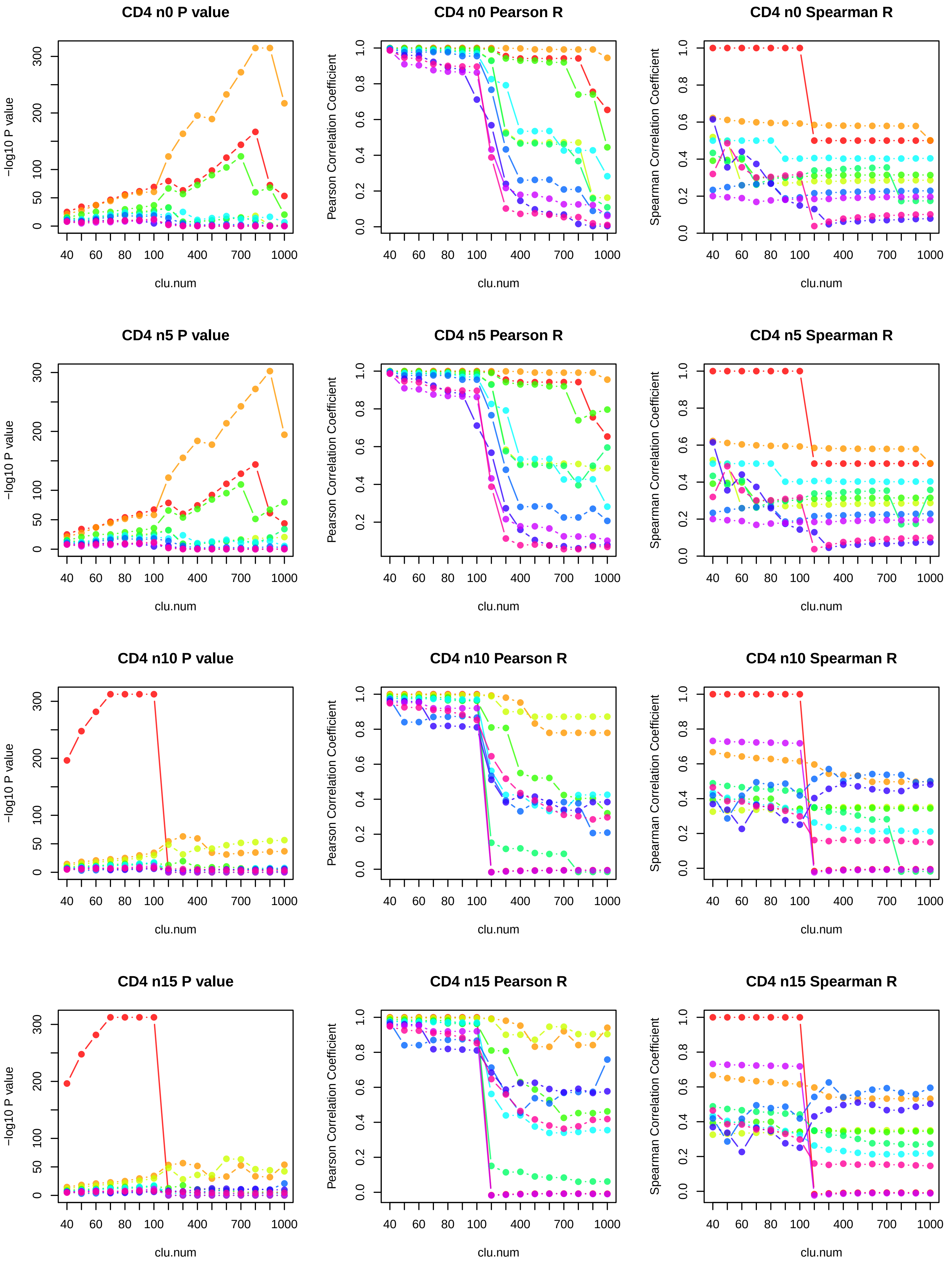

### Supplemental Figure 6

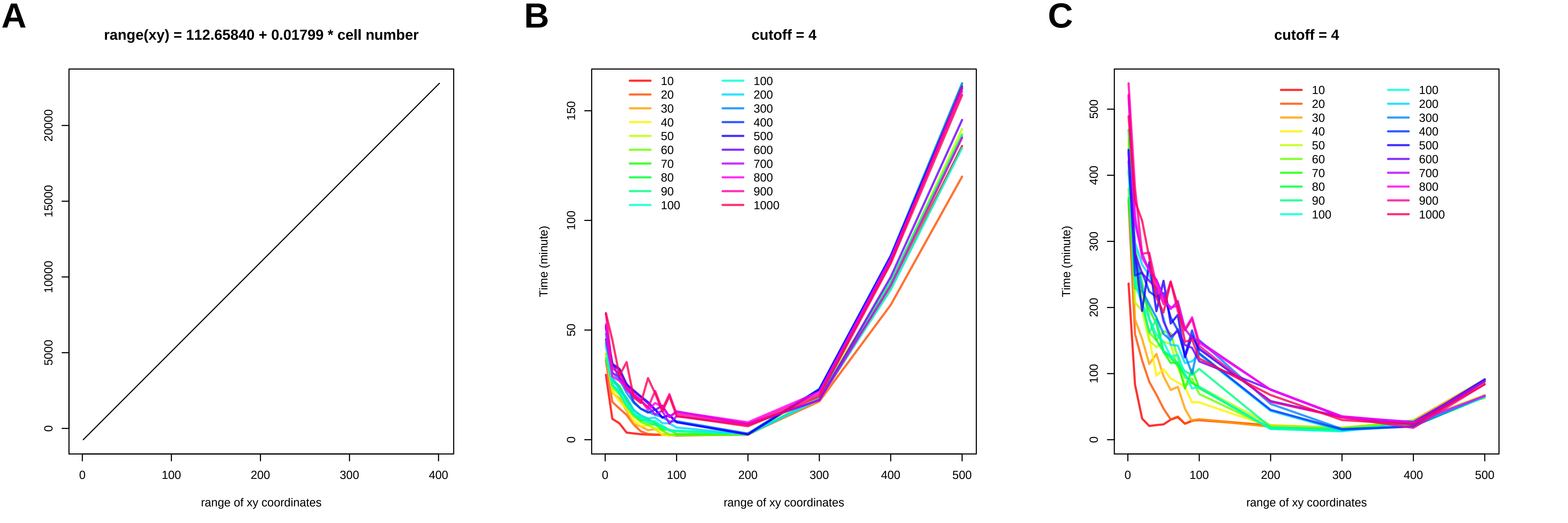
